# Suboptimal larval habitats modulate oviposition of the malaria vector mosquito *Anopheles gambiae*

**DOI:** 10.1101/027367

**Authors:** Eunho Suh, Dong-Hwan Choe, Ahmed M. Saveer, Laurence J. Zwiebel

## Abstract

Selection of oviposition sites by gravid females is a critical behavioural preference in the reproductive cycle of *Anopheles gambiae*, the principal Afrotropical malaria vector mosquito. Several studies suggest this decision is mediated by semiochemicals associated with potential oviposition sites. To better understand the chemosensory basis of this behaviour and identify compounds that can modulate oviposition, we examined the generally held hypothesis that suboptimal larval habitats give rise to semiochemicals that negatively influence the oviposition preference of gravid females. Dual-choice bioassays indicated that oviposition sites conditioned in this manner do indeed foster significant and concentration dependent aversive effects on the oviposition site selection of gravid females. Headspace analyses derived from aversive habitats consistently noted the presence of dimethyl disulphide (DMDS), dimethyl trisulphide (DMTS) and 6-methyl-5-hepten-2-one (sulcatone) each of which unitarily affected *An. gambiae* oviposition preference. Electrophysiological assays across the antennae, maxillary palp, and labellum of gravid *An. gambiae* revealed differential responses to these semiochemicals. Taken together, these findings validate the hypothesis in question and suggest that suboptimal environments for *An. gambiae* larval development results in the release of DMDS, DMTS and sulcatone that impact the response valence of gravid females to directly modulate the chemical ecology of oviposition site selection.

## Introduction

Mosquito-borne malaria remains among the greatest threats to global human health ^1^. Inasmuch as effective vaccines are still elusive, the widespread use of the current set of anti-malarials and insecticides has contributed to the rise in resistance to these agents in both pathogens and vectors, respectively ^2,3^. In this light, vector control remains among the most effective methods in reducing disease transmission ^4^.

A critical feature of improved vector control programs is an enhanced understanding of the mechanistic basis of both vector competence and vectorial capacity. Together with blood-meal host selection/preference, the search for oviposition sites representing optimal larval breeding habitats are crucial decisions in mosquito reproductive cycles that directly impact vector population size and, accordingly, vectorial capacity ^5^. In the course of oviposition site selection in the field gravid females dynamically process multiple signals including hygroscopic, olfactory, tactile, thermal or visual cues to assess larval breeding sites ^6-8^. In particular, semiochemicals derived from multiple sources that include mosquito, microbiological, predator and plant-derived constituents that comprise the ecosystems of immature mosquitoes play critical roles in oviposition site selection ^6-8^.

In the field, gravid *An. gambiae* which is acknowledged to represent several distinct and newly emerging species including *An. coluzzii* and *An. amharicus* ^9^ oviposit directly on a diverse spectrum of habitat water as well as surrounding muds where larvae hatch and find their way to nearby water ^10^. Accordingly, immature *An. gambiae* occurs in aquatic breeding sites with diverse biological and physico-chemical characteristics ^11-13^. In addition to abiotic factors such as water vapour, hydration, or visual contrast of oviposition sites ^14-16^, studies have suggested chemical signals derived from biotic components of breeding sites such as microbial larval food sources ^17-19^ or predators/competitors ^20-22^ influence the oviposition behaviour of gravid *An. gambiae*. Indeed, oviposition site selection behaviour of mosquitoes has been postulated to evolve toward maximization of offspring fitness interacting with multiple biotic factors ^23-27^. Consistent with this hypothesis, oviposition behaviour of gravid female mosquitoes is associated with conspecific larval density ^26-28^, suggesting that population size of immature mosquitoes are maintained at near optimal levels by selective oviposition of gravid females utilizing unknown cues associated with the pre-existing larval population that reflects the expected fitness of those populations. Specifically, presence of conspecifics in low density could be an indication of suitable breeding sites (e.g., appropriate larval food level along with the absence of predators) where oviposition activity tends to be stimulated/increased, whereas increased larval competition for limited resource result in overall reduction in the fitness of progenies where the oviposition activity tends to be deterred/reduced ^23,24,29,30^.

In order to further characterize *An. gambiae* aversive oviposition cues specifically associated with suboptimal rearing conditions, we generated artificial habitats containing overcrowded and resource-deprived larvae that would be expected to have a repellent effect on ovipositing females. To examine this, we developed a two-choice oviposition bioassay to evaluate the oviposition preferences of gravid females to pre-conditioned larval water (LW) and characterized its major volatile constituents in both behavioural and electrophysiological paradigms with gravid adult females. Our studies validate the hypothesis that suboptimal larval habitats repel gravid females and identify dimethyl disulphide (DMDS), dimethyl trisulphide (DMTS), and 6-methyl-5-hepten-2-one (sulcatone) as a specific semiochemicals involved in modulating the oviposition behaviour of *An. gambiae*.

## Results

### Oviposition behaviour of *An. gambiae* is mediated by larval water-derived volatiles

A series of oviposition assays were conducted to examine the effects of high larval density coupled with limited nutrient resources on the oviposition behaviour of *An. gambiae* gravid females. Here, studies utilized a range of LW treatments that varied the duration of time without food and larval density to establish conditions that would elicit the strongest effects on olfactory-driven oviposition behaviours of gravid females.

In the first experiment, preconditioned LW samples were obtained by varying number of late instars (5, 10, 50, 100, 300 larvae) incubated for 72 hours and were used in dual choice oviposition bioassays between egg laying cups containing either LW (treatment) or control water (CW, untreated). In these studies, significant aversive responses (as indicated by negative oviposition index [OI] values) that reduced oviposition increased relative to the number of larvae in LW treatments with OI values ranging from −0.33 to −0.77 (50 to 300 larvae, Fig. 1B). Lower larval densities (5 and 10 larvae) resulted in LW samples with no effect on the oviposition preference of gravid females (Fig. 1B). The number of surviving larvae was counted to estimate the larval survivorship for each LW treatment. The survival of larvae was significantly reduced (54%) at higher larval densities (300 larvae) while lower densities (5, 10, 50, 100 larvae) had similar larval survival ranging from 94% to 100% (Fig. S1A). Importantly, the number of total eggs collected from two oviposition cups in the dual choice bioassay was not affected by the initial number of larvae used for LW conditioning (ANOVA, *F_5,122_* = 1.16, *p* = 0.33; Fig. S2A) suggesting the presence of compounds in the assay cages neither stimulated nor deterred oviposition of gravid females. We next varied incubation time (0 h, 24 h, 48 h, 72 h) of 300 larvae to generate LW samples. Significant aversion was observed in response to LW generated when larvae were held beyond 24 h with OI values ranging from −0.47 to −0.76 while no effect was observed for CW treatments (Fig. 1C), indicating the repellent effect of LW on gravid females is positively correlated with both the initial larval density and the duration of LW conditioning. In these studies, larval survival was significantly reduced over time ranging from 94% to 52% and increasing numbers of dead larvae (often with lost body parts) were observed via visual inspections (Fig. S1B and Fig. S1C). Once again, the overall number of total eggs collected from the two oviposition cups in the dual choice assay was not affected by duration of the LW conditioning (ANOVA, *F_3,77_* = 1.34, *p* = 0.27; Fig. S2B). In addition, LW samples with increased larval densities, varied conditioning time or larval age showed no additional increase in OI value (Fig. S3) as compared to maximum OIs observed in the previous bioassays (Fig. 1B and Fig. 1C). Thus, together with the results that the LW samples generated from 300 larvae incubated for 72 hours exhibited a significantly increased larval mortality in the survival analyses (Fig. S1A and Fig. S1B), LW samples generated by this condition used as standard LW hereafter.

**Figure 1.**
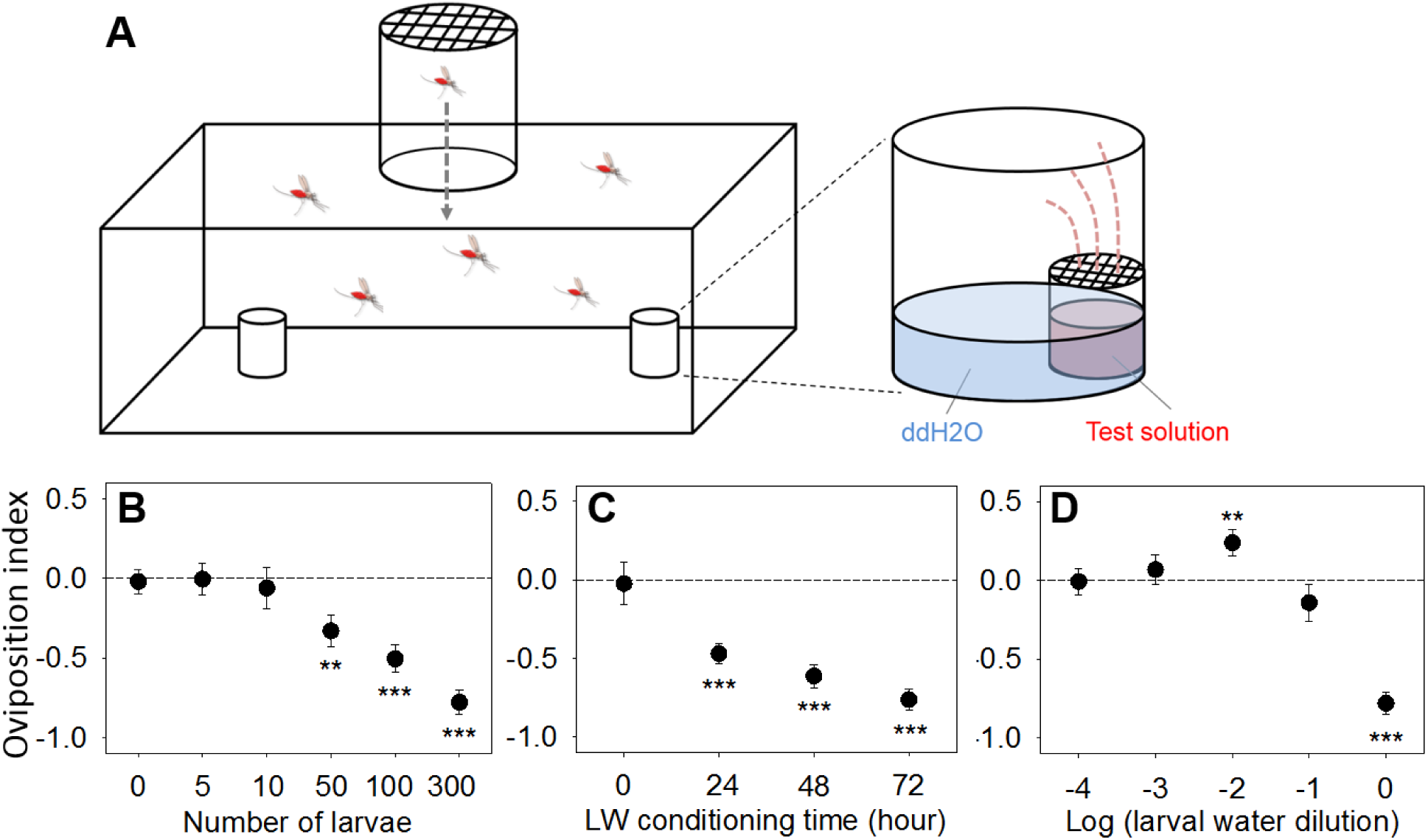
Behavioural response of *An. gambiae* gravid females in oviposition dual choice assay between control water and test solution. (A) Schematic of behavioural assay designed to examine olfactory-driven responses (see Methods for details). Oviposition preference of gravid females to larval water samples with varied treatment by (B) incubating different number of late instars for 72 h, (C) incubating 300 late instars for different time period, and (D) diluting LW sample obtained by incubating 300 late instars for 72 h. Asterisks represent significant OI values different from zero (***, *p* < 0.001; **, *p* < 0.01; Wilcoxon signed-rank test, two-sided). Error bar = s.e.m. (n = 17 ~ 36).

This concentration dependent effect was also examined by observing the behavioural responses elicited by serial dilutions (in ddH2O) of the most active LW sample (i.e., standard LW) in dual choice oviposition bioassays. Consistent with prior results, significant aversion was observed in oviposition bioassays using undiluted LW (OI = − 0.78 ± 0.071; mean ± s.e.m; n = 18; Wilcoxon signed-rank, *Z* = −84, *p* < 0.0001) although this effect was not observed when LW was diluted as low as 10 fold (Fig. 1D). Interestingly, when LW samples were further diluted to 100 fold, a significant degree of attraction was observed (OI = 0.23 ± 0.11; n = 17; Wilcoxon signed-rank, *Z* = 57; *p* < 0.01), and once again, this effect decreased as the concentration of LW was further reduced (Fig. 1D). As in previous assay, the total number of *An. gambiae* eggs collected from the two oviposition cups was not affected by concentration of conditioned LW (ANOVA *F_4,92_* = 0.73, *p* = 0.58; Fig. S2C).

### Oviposition behaviour of *An. gambiae* is affected by larval water specific unitary compounds

Headspace analyses of LW samples were carried out using gas chromatography (GC) - mass spectrometry (MS) with solid phase micro-extraction (SPME) fibres. In addition to several trace compounds, significant concentrations of dimethyl disulphide (DMDS), dimethyl trisulphide (DMTS) and 6-methyl-5-hepten-2-one (sulcatone) were consistently detected in the headspace of LW samples, but not in the headspace of CW samples (Fig. 2). Quantitative analyses of the headspace of standard solutions containing known amounts of individual compound indicated that 10^−7^ M DMDS, 10^−8^ M DMTS and 10^−8^ M sulcatone in aqueous preparations produced the peak integration values that were most similar to those observed in LW headspace (i.e., within a range of one order of magnitude) (Fig. 2). Based on the ratio between peak integration values from headspaces derived from standard compounds at those concentrations and those integration values derived from the corresponding peaks from six LW headspace samples, (see Methods for detailed information) the approximate concentrations of DMDS, DMTS and sulcatone in LW were estimated to be (1.72 ± 0.47) × 10^−7^, (3.55 ± 0.49) × 10^−9^, and (2.75 ± 1.07) × 10^−9^ M (mean ± s.e.m; n = 6), respectively.

**Figure 2.**
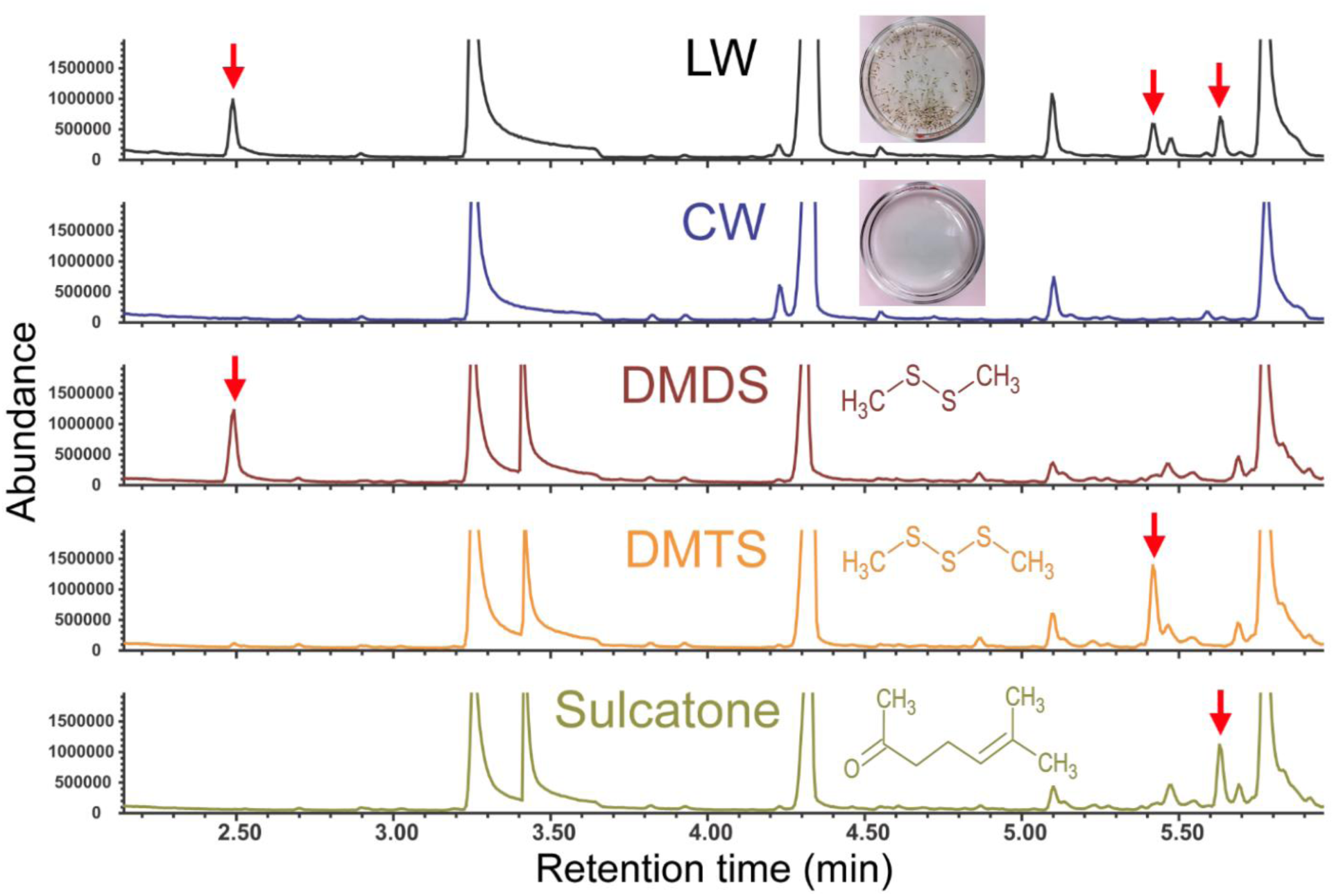
SPME analysis of headspace volatiles of water samples. Partial chromatograms are shown for volatile samples taken from larval water (LW), control water (CW), and standard DMDS (10^−7^ M), DMTS S (10^−8^ M) and sulcatone (10^−8^ M). Peaks for DMDS, DMTS and sulcatone are marked with red arrows. No additional LW specific compounds were detected beyond the retention time of sulcatone. Large peaks with retention times of approximately 3.2, 4.3 and 5.8 min represent impurities possibly introduced during sample preparations and/or chemical analyses, which are present in all samples including LW and CW.

The behavioural responses of gravid female *An. gambiae* to egg laying cups containing DMDS, DMTS and sulcatone over various concentrations was examined in dual choice oviposition bioassays. In these studies, egg laying cups with 10^−8^ M and 10^−7^ M DMDS elicited aversive response with OI value of 0.25 ± 0.08 (mean ± s.e.m.; n = 32; Wilcoxon signed-rank, *Z* = −152, *p* < 0.01) and −0.15 ± 0.066 (n = 26; Wilcoxon signed-rank, *Z* = −77, *p* < 0.05) respectively, while moderate attraction was observed at 10^−9^ M DMDS (OI = 0.16 ± 0.082; n = 35; Wilcoxon signed-rank, *Z* = 122, *p* < 0.05) (Fig. 3). Unlike DMDS, DMTS and sulcatone did not show significant attractancy at any concentration but instead elicited relatively increased aversive responses in similar dose dependent manners. Here, 10^−8^ M and 10^−7^ M DMTS had significant repellency with OI values of −0.43 ± 0.11 (n = 15; Wilcoxon signed-rank, *Z* = −53, *p* < 0.01) and −0.25 ± 0.089 (n = 17; Wilcoxon signed-rank, *Z* = −50, *p* < 0.05), respectively. Similarly, sulcatone elicited significant aversive responses at 10^−6^ M (OI = −0.4 ± 0.11; mean ± s.e.m.; n = 14; Wilcoxon signed-rank, *Z* = −43, *p* < 0.01) while the OI value at 10^−5^ M sulcatone approached significance (−0.26 ± 0.12; n = 17; Wilcoxon signed-rank, *Z* = −37, *p* = 0.081) (Fig. 3). The total number of eggs collected from both oviposition cups was not affected by differing concentrations of DMDS, DMTS and sulcatone (ANOVA, *p* > 0.05; Fig. S2).

**Figure 3.**
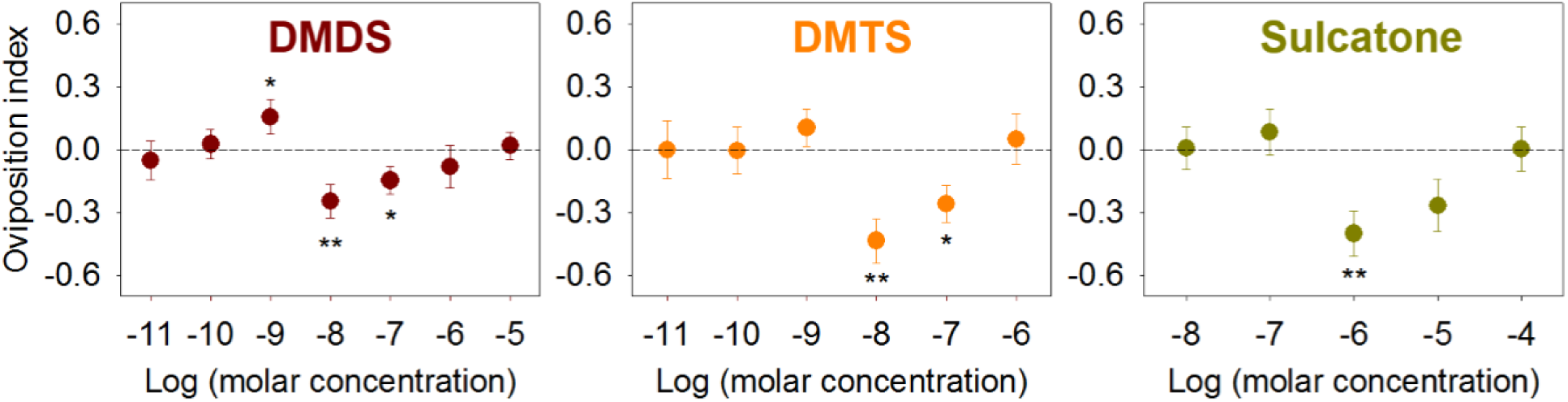
Behavioural response of *An. gambiae* gravid females in oviposition dual choice assay between control water and DMDS, DMTS and sulcatone with serial dilutions. Asterisks represent significant OI value different from zero (**, *p* < 0.01; *, *p* < 0.05; Wilcoxon signed-rank test, two-sided). Error bar = s.e.m. (n = 14 ~ 35).

### The antennae, maxillary palp and labellum of gravid mosquitoes respond to larval water, dimethyl disulphide, dimethyl trisulphide and sulcatone

In an effort to further explore the basis of DMDS, DMTS and sulcatone on *An. gambiae* oviposition preferences we examined the electrophysiological responses of antennae, maxillary palp and labellum in gravid mosquitoes. These responses were initially characterized using electroantennogram (EAG), electropalpogram (EPG) and electrolabellogram (ELG) assays that measure transcuticular voltage changes derived from collective neuronal responses ^31^. In these studies, each chemosensory appendage exhibited significant, albeit differential, sensitivity to individual stimuli as well as to complex LW samples. Specifically, the maxillary palp maintained a significant response to a 10 fold LW dilution while antennal and labellar responses were restricted to only undiluted LW (Fig. S4). Similarly, the antennae of gravid *An. gambiae* was considerably more sensitive to sulcatone with significant responses at 10^−5^ M as compared to DMDS or DMTS where thresholds were 10^−2^ M and 10^−3^ M, respectively (Fig. 4A). The maxillary palp was somewhat more sensitive to DMDS and DMTS significantly responding as low as 10^−4^ M and 10^−5^ M, respectively, with significant sulcatone responses as low as 10^−4^ M (Fig. 4A). The labellum was more sensitive to sulcatone and DMTS relative to DMDS with significant responses as low as 10^−4^ M, respectively (Fig. 4A). Beyond those threshold levels, the chemosensory appendages exhibited odour-dependent polarization showing diverse response traces. While all three compounds elicited downward (depolarization) responses in EAGs studies, upward (hyperpolarization) responses were frequently observed in EPG and/or ELG studies (Fig. 4A and Fig. S5A).

**Figure 4.**
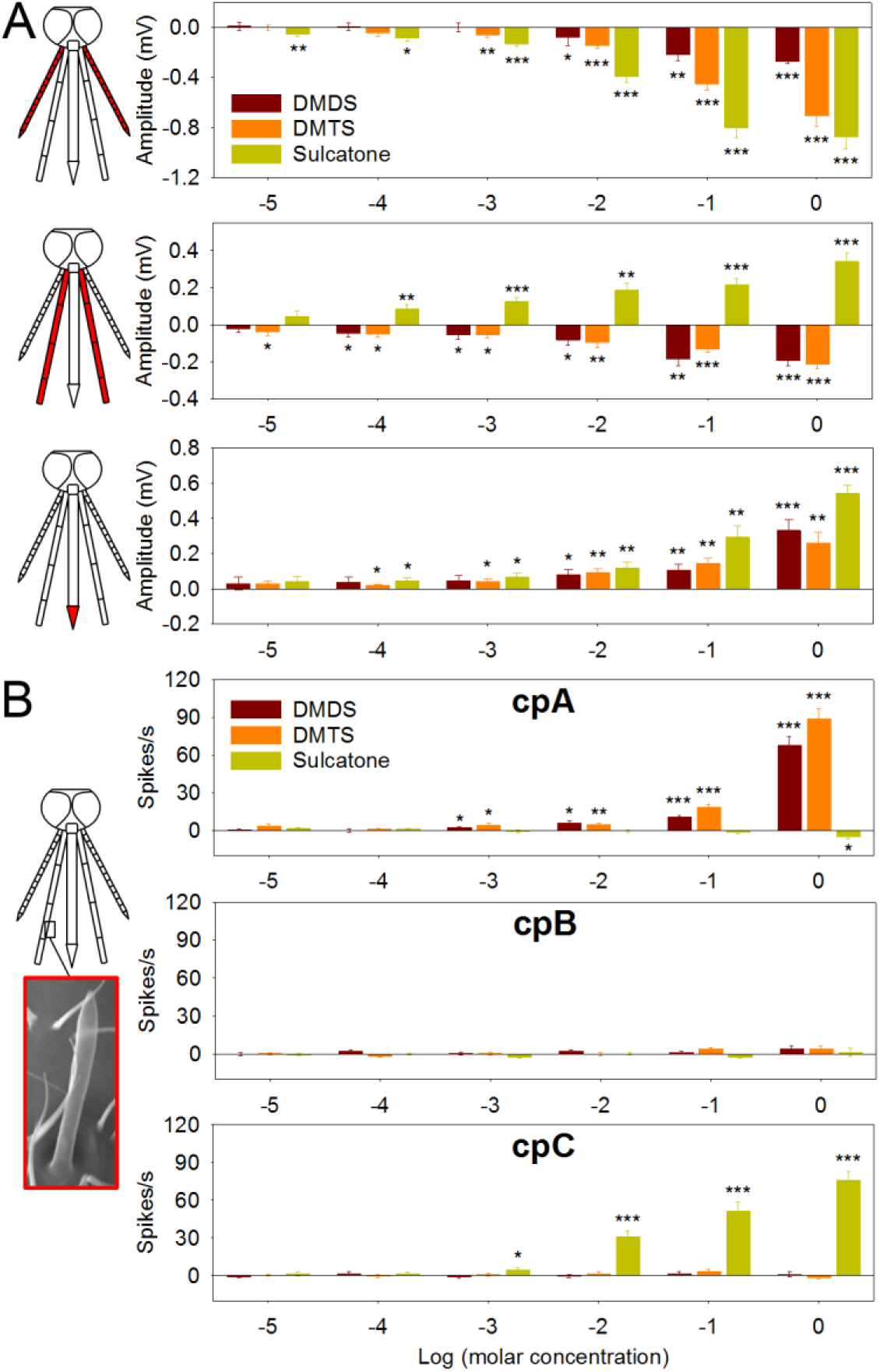
Relative electrophysiological response of *An. gambiae* females to DMDS, DMTS and sulcatone ne. Electrophysiological responses of (A) antenna, maxillary palp and labellum (top to bottom; each chemosensory organ is highlighted in red in a schematic diagram of mosquito head) and (B) neurons in capitate peg sensillum (highlighted in a red box; picture modified from ^32^) of maxillary palp, expressed as response difference to solvent control (oil). Y axis represents response amplitude subtracted by control values and X axis represents log transformed molar concentration. Asterisks represent significant response amplitude different from zero (***, *p* < 0.001; **, *p* < 0.01; *, *p* < 0.05; one sample *t*-test, one-sided). Error bar = s.e.m. (n = 7~10).

### Neurons in capitate peg sensillum of the maxillary palp respond to dimethyl disulphide, dimethyl trisulphide and sulcatone

We next utilized single sensillum electrophysiological recordings (SSR) to provide a detailed characterization of the neuronal responses to DMDS, DMTS and sulcatone on the maxillary palp of gravid *An. gambiae*. Here, well characterized capitate peg (cp) sensilla containing triad chemosensory neuron subpopulations (cpA, cpB and cpC) are uniformly distributed ^32^. In these studies, we consistently observed biphasic responses such that stimulation with DMDS and DMTS initially evoked excitatory increases in the CO_2_ sensing cpA neuron spike frequency. This was followed by discrete inhibitory responses of 0.2 ~ 0.3 sec in which cpA spike activity was reduced before the recovery of spontaneous (base line) neuronal activity (Fig. S5B). Unlike DMDS or DMTS, sulcatone elicited a prolonged cpC odorant receptor neuron (ORN) excitatory response for several seconds after the onset of stimulus while cpA ORN responses were inhibited for more than one second (Fig. 4B and Fig. S5B). Spike sorting analyses of these traces revealed dose-dependent responses for cpA neurons which showed significantly increased spike frequencies in response to as low as 10^-3^ M DMDS and DMTS. In contrast, cpB and cpC ORNs did not respond to DMDS nor DMTS (Fig. 4B). Unlike DMDS or DMTS, sulcatone elicited significantly increased spike frequencies for cpC neurons in a dose dependent manner down to 10^-3^ M while cpA and cpB neurons displayed no response to sulcatone except that the cpA neuron was significantly inhibited by 1 M sulcatone (Fig. 4B).

## Discussion

Despite many studies that posit the role of chemical signals in mediating oviposition site selection of *An. gambiae*, a paucity of validated (in either laboratory or field studies) oviposition-related semiochemicals have been characterized in this medically important mosquito. Indeed, at this point and apart from the general role of water vapour as an attractant ^16^ three volatile, unitary oviposition cues have been identified in *An. gambiae*: 2-propylphenol, 4-methylcyclohexanol and cedrol ^33,34^. We now report studies that utilize a laboratory-based oviposition bioassay specifically designed to examine olfactory driven behavioural responses of ovipositing females to test the generally held hypothesis that unsuitable conditions for larval development results in the release of semiochemicals that repel gravid female mosquitoes in search of oviposition sites. Taken together, our data support the hypothesis and show that potential oviposition sites conditioned by larvae maintained under poor conditions are actively avoided by gravid female mosquitoes. Consistent with prior studies ^28,35^, the degree of repellency was dependent on conditioning characteristics such as duration and larval density (Fig. 1). Repellency appears to arise strictly as a result of overcrowding as increases in larval density (e.g. from 10 to 50/dish) significantly induced a repellent effect (Fig. 1B) without affecting larval survival (Fig. S1A). Furthermore, LW-derived repellency increased along with incubation time (Fig. 1C) coinciding with a decrease in the larval survival rate from 94% to 52% (Fig. S1B). This suggests that increased incubation time and the density of deceased larval remains may additively increase to the repellent effect of LW. Identification of the precise source(s) of volatile compounds should provide useful information whether the repellency of LW is due to production of volatiles by larval starvation/crowding effects or through increase in the concentration of volatiles generated by other constituents in LW (e.g., micro-organisms, etc.).

While the inability of conditioned LW samples produced at lower larval densities to attract gravid females (Fig. 1B) is inconsistent with the “dual effect” mosquito oviposition regulation model that balances both attraction and repulsion ^27^, similar dual effects were observed using serial dilutions of LW samples that elicited the strongest aversive effects (Fig. 1D). This suggests that increased larval density may generate not only quantitatively but also qualitatively different odour profiles compared to those derived from LW generated at lower larval densities. It is possible that in the field gravid female *An. gambiae* are more attracted to larval habitats previously occupied to a large number of conspecific larvae and subsequently replenished with fresh water while virgin or under-populated oviposition sites are relatively neutral. This hypothesis is also supported by a previous study in which attraction was frequently observed in LW samples with low number of larvae with field collected water from natural larval habitats while neutral effects or modest repellency was observed from LW samples using only distilled water ^28^.

While we cannot, as yet, define the precise source of these chemical signals which would require comprehensive chemical analyses on different larval water treatments as well as identification of LW constituents such as micro-organisms, it is evident that our laboratory-based LW conditioning paradigm induces high larval stress and mortality that likely release and/or promote the accumulation of a range of potential semiochemicals. In addition, this process is likely to liberate a diverse microbiological population derived from damaged, decayed or predated larvae that also constitutes a potential source of semiochemicals. Indeed, gravid *An. gambiae* was significantly attracted by volatiles emitted by several bacteria species from larval midgut or natural larval habitats ^18^. In particular, *Stenotrophomonas maltophilia* bacteria isolated from natural larval habitats has been identified to repel gravid *An. gambiae*^19^. Similarly, laboratory-based oviposition of *Ae. aegypti* is also affected by bacterial volatiles in dose dependent manners ^36,37^. Overall, in addition to diverse effects of a large number of non-olfactory factors (e.g., tactile, visual cues, etc.) as observed in previous studies ^14,15,38^, oviposition in *An. gambiae* is likely to also be associated with the complex chemical ecology of native oviposition sites together with the interactive effects derived from the population dynamics of microorganisms and pre-adult stages of the mosquito.

We have identified DMDS, DMTS and sulcatone as significant and specific volatile components of the laboratory-derived LW semiochemical blend that has dose-dependent oviposition effects on gravid female *An. gambiae*. These compounds are well established semiochemicals across insect taxa including host seeking hematophagous insects 39-44, and sulcatone is often considered as a repellent or “masking” odorant in mosquitoes 45,46. While oviposition data presented here is consistent with that mode of action, we cannot conclude these compounds either as unitary compounds or in blend, are necessary and sufficient for the complete spectrum of repellent effects of LW.

Estimated LW source concentrations of DMDS, DMTS and sulcatone ranged from 10^−9^ M to 10^−7^ M, which was overlapping or close to the bioactive concentrations that showed repellent effects in the behavioural assays of unitary compounds (10^−8^ M to 10^−6^ M; Fig. 3). While additional studies are required to fully understand the contribution of each of these compounds to the repellent effects of LW (e.g., source concentrations, temporal emission dynamics etc.), the discrepancy between our estimated LW source concentration and the actual bioactive concentrations also reflect the inherent differences between chemical analyses and behavioural bioassays. During the chemical analyses, the extraction devices (i.e., SPME fibre in a sealed sample vial) were exposed in a static headspace within the vial in which the volatiles were equilibrated between the gas phase and the liquid phase, producing a partial pressure. In contrast, during the behavioural bioassays, the test organisms (i.e., gravid female in the oviposition arena) were exposed to the volatiles that were not contained in a sealed vial, perhaps experiencing a freely diffusing concentration gradient within an odour plume. These may in part explain the discrepancy between the estimated LW source concentration of sulcatone (~10^−9^ M) and its oviposition bioassay active concentration (~10^−6^ M) (Fig. 3). Furthermore, inasmuch as oviposition could occur anytime during the overnight bioassay ^47^, this behaviour is likely to be affected by distinct headspace volatiles as they are continuously being emitted from egg cups, an effect which could contribute to the narrow responsive range of bioactive concentrations resulting in sudden shifts in oviposition preference (Fig. 3) as often observed in prior studies ^36,48,49^. The estimated concentrations would, therefore, represent only approximate abundances of these compounds in LW, and absence of significant effects of tertiary mixture of DMDS, DMTS and sulcatone at the estimated concentrations in behavioural assays (data not shown) is not surprising considering the potential limitation in concentration estimations apart from the difficulty to precisely mimic complex nature of LW where vaporization dynamics of individual odours during oviposition assay periods are unknown.

We observed significant transcuticular electrophysiological responses to complex volatiles emitted from undiluted pre-conditioned LW as well as unitary DMDS, DMTS, sulcatone from the antenna, maxillary palp, and labellum of gravid *An. gambiae* (Fig. 4A). It is noteworthy that EAG responses to natural larval habitats have previously been reported in *An. gambiae* suggesting that volatiles from those habitats may affect oviposition behaviour of gravid females ^50^. Using EAG/EPG/ELG recordings we observed concentration dependent responses of gravid females to the three LW specific compounds with differential sensitivity for each chemosensory organ. This suggests that modulation of oviposition behaviours in *An. gambiae* is associated with multiple chemosensory organs differentially responding to each compound while, at this point, the relative contribution of each chemosensory appendage or cell type in mediating *An. gambiae* oviposition behaviours has yet to be precisely mapped.

Interestingly, significant responses of gravid females to the three compounds were observed at relatively lower concentrations (e.g., 10^−9^ to 10^−6^ M) in dual choice behavioural assays while electrophysiological responses were limited to higher concentrations (e.g., DMDS or DMTS) (Fig. 3 and Fig. 4). These results suggest oviposition behaviours could be elicited by continuous exposure of volatiles at concentrations that are not necessarily EAG/EPG/ELG positive. Electrophysiology studies which measure response profiles of individual appendages or sensory neurons are presumed to display a significantly reduced response sensitivity relative to the intact insect where evolution has generated a highly efficient biological platform (i.e. the central nervous system and other networks) to integrate, filter and otherwise process numerous signals from peripheral sensory systems to generate behavioural outputs^51^. For these reasons, as well as a range of technical considerations, some of which were discussed above, responsive thresholds observed in electrophysiological studies are not necessarily expected to reflect the behaviourally active semiochemical concentrations established in organismal bioassays.

While a comprehensive characterization of antennae and labellum SSR responses is challenging for *Anopheles* due to the complexity in sensillar type and location, the maxillary palp provides an ideal model. This sensory appendage expresses a considerably more narrow receptor repertoire across a uniform population of capitate peg sensillum containing only three (cpA, cpB and cpC) sensory neurons^32^. DMDS and DMTS induced excitatory responses as well as post-stimulation inhibition in the cpA neuron of gravid *An. gambiae* that is devoid *AgOr* (*Anopheles gambiae* odorant receptor) of expression (Fig. 4B and Fig. S5B) ^32^. These cpA responses were significantly more sensitive at lower concentrations of DMDS and DMTS while both the *AgOr* expressing cpB or cpC neurons were not responsive (Fig. 4B and Fig. S5B). These data implicate the cpA neuron as the major sensing neuron for DMDS and DMTS on the maxillary palp of *An. gambiae*. While devoid of *AgOrs*, the cpA neuron is known to express the gustatory receptors *AgGr22, 23, 24* that are principally involved in responses to CO_2_ and have been shown to display sensitivity to a range of other semiochemicals ^32,52^. In contrast, sulcatone induced dose dependent, excitatory responses in cpC neurons, implicating *AgOr*28 as a putative oviposition receptor on the maxillary palp of *An. gambiae* females (Fig. 4B and Fig. S5B). The cpC and cpB responses are discriminated based on amplitude differences and the shape of spikes (Fig. S5B). Interestingly, dose dependent cpC tonic responses of up to 10 seconds (Fig. S5B) were observed while cpA neurons were significantly inhibited for ~one second after high-concentration stimulation with 1 M sulcatone (Fig. 4B and Fig. S5B). The inhibition of cpA neurons with the simultaneous excitatory response of cpC neurons is possibly due to ephaptic coupling between these neurons^53^.

At the molecular level, a large number of sulcatone-tuned *An. gambiae* odorant receptors (AgORs) expressed across all three chemosensory appendages are likely to be associated with the sulcatone responses of gravid *An. gambiae* females ^54,55^. While DMTS receptors have not been molecularly identified, the distinct cpA-centred neuronal response among maxillary palp neurons elicited by this compound and DMDS suggest they are encoded by either the gustatory or ionotropic receptors expressed in those non-AgOR expressing cells on the maxillary palp of *An. gambiae* ^32,52,56^. Elucidation of the precise relationships between molecular receptors and these oviposition behaviours will be an important component in establishing a path for the development of vector control strategies that target this critical step in the reproductive lifecycle of *An. gambiae*.

This study provides evidence that olfactory driven oviposition behaviours are modulated by volatiles associated with suboptimal larval breeding sites. Specifically, starvation and/or over-crowding of larvae increased the emission of volatile semiochemicals that elicited aversive effects on ovipositing gravid females whereas diluted LW also elicited attractant effects in keeping with a widely accepted mosquito oviposition regulation model ^27^. We have identified DMDS, DMTS and sulcatone as distinct, behaviourally active components of this response that elicit dose-dependent attractant and/or repellent effects, and propose these compounds are associated with regulating oviposition behaviour of *An. gambiae*. How the olfactory components revealed in this study fit the complex dynamics of oviposition biology of *An. gambiae* in natural populations under field conditions should be addressed in the future studies.

## Materials and Methods

### Mosquito rearing

*Anopheles coluzzii* (SUA 2La/2La), originating from Suakoko, Liberia, was maintained in the Vanderbilt Insectary. Mosquito rearing primarily followed a lab protocol developed in a previous study ^33^. In brief, larvae were reared under standardized conditions (> 2 cm^2^ water surface per larva with 1 litre of dH2O per larval pan) by adding food *ad libitum* in environmental chamber (27°C, 80% relative humidity, light:dark = 12:12 h) to ensure consistent larval development and large size adults. Pupae were collected and eclosed in a cage, and females and males were allowed to mate with constant access to 10% sucrose solution. For blood feeding, 6 to 7 days old females were provided with human blood (BioChemed, Winchester, VA) by using Hemotek membrane feeding system (Hemotek, Lancaster, UK), and 4.5% CO2 was utilized to promote blood feeding.

### Preparation of conditioned larval water (LW)

All LW treatment used larvae reared under standard rearing condition described above. Late 3^rd^ or early 4^th^ instars were collected in dH_2_O and washed/filtered several times through a wire sieve and then transferred into 50 ml borosilicate glass bottle with 20 ml of HPLC grade dH_2_O for incubation at 27°C. Conditions used here for LW treatment were comparable to a previous study ^28^. Control water (CW) was prepared using HPLC grade dH2O without larvae. Larvae were removed from LW using a metal sieve (standard sieve No. 40; opening size = 0.420 mm) after completion of larval incubation. In order to minimize contamination of random odours in chemical analyses, materials used for LW and CW sample preparations (i.e., glass containers) were first cleaned with detergent, and then serially washed with distilled water, methanol and finally methylene chloride, and cleaned materials were incubated at 75°C overnight followed by complete desiccation before use. LW and CW samples were kept at 4°C for no more than one week before being used for the oviposition assay and headspace chemical analysis.

### Dual choice oviposition assay

Laboratory based oviposition bioassays were conducted in a growth chamber following the previously established protocol ^33^. To prepare gravid female mosquitoes, 6 to 7 days old females were blood fed, and fully engorged females were transferred to a separate cage with constant access to 10% sucrose solution. Preparation of the experimental setup began around 1 hour prior to scotophase in the environmental chamber under the same conditions used for larval rearing as described above. Two days after blood feeding, 10 gravid females were transferred into the “releasing chamber” with a screen on top, and the females were allowed to enter the assay cage (polypropylene, length = 37.8 cm, height = 15.2 cm, width = 13 cm) through a centre pathway (diameter = 6 cm) after dark cycle began (see Fig. 1A). A borosilicate glass vial (capacity = ~ 1 ml, 14.65 × 19 mm; Qorpak, Bridgeville, PA) with a screen on top (10 × 10 mm) was used to contain 1 ml of test or control water placed in order to exclude the effect of tactile cue on the oviposition behaviour of gravid females. The vials were placed inside of the egg cups (PET, top diameter = 4.5 cm, height = 4.1 cm, bottom diameter = 2.9 cm) farther from the releasing chamber with their sides in contact with the inner side of the egg cups. The two egg cups (control and test) were 26 cm apart. The egg cups were filled with 7 ml of ddH_2_O as an oviposition substrate. In this experimental design, gravid females were not allowed to touch aqueous solution within the vial, thus oviposition preference observed in the bioassays was assumed to be driven by olfactory cues. Location of egg cup containing test water was rotated between assay cages, and the assay cages were randomly placed within a larger enclosure (acrylic, 86 × 120 × 86 cm) to minimize external effect on the oviposition preference. For the preparation of test waters of lower concentrations, LW were diluted in ddH2O, and DMDS, DMTS and sulcatone was dissolved and diluted in 0.1% DMSO as standard solutions. DMDS, DMTS, sulcatone and DMSO (≥ 98% purity) were obtained from Sigma-Aldrich, Inc. All assay cages were cleaned by using 70% EtOH and fully dried before each assay. Gravid females were allowed to oviposit during scotophase, and collected eggs were counted in the following morning. All bioassays were conducted using at least three different batches of mosquitoes.

The total number of collected eggs were used to calculate oviposition index (OI) using formula OI = (N_t_ –N_c_)/(N_t_+N_c_) ^57^ with N_t_ = number of eggs collected in the egg cup with LW or test volatiles (i.e., larval conditioned water or test compound in the vial), and N_c_ = number of eggs collected in the egg cup with CW volatiles. Oviposition preference of gravid females was determined by OI values using Wilcoxon signed rank test (*p* = 0.05, two-sided; JMP 8.0.1; SAS Institute, Cary, NC), and the nonparametric method was used due to non-normality of data. If the OI values were significantly different from zero with positive or negative values, the subject was considered to have attractant or repellent effect on oviposition behaviour of gravid females, respectively. In order to examine any stimulatory or deterrent effect of test water/compounds on the oviposition of gravid females, ANOVA (*p* = 0.05; JMP 8.0.1; SAS Institute, Cary, NC) was used to test the effect of treatment variables (e.g., larval density, incubation time, concentration, etc.) on the total number of eggs collected in a cage. If there’s no effect of treatment variable, we considered oviposition of gravid females were neither stimulated nor deterred by the presence of test water/compounds. Cages that collected less than 200 eggs were excluded from all analysis, and 74.9 ± 3.7% (mean ± s.e.m.; n = 6) of raw data obtained from each set of behavioural assay (see Fig. 1B, Fig. 1C, Fig.1D and Fig. 3 for replication numbers for each set of behavioural assay) were used.

### Head space analysis on pre-conditioned larval water

Headspace volatiles of LW samples (300 early 3^rd^ or late 4^th^ instars incubated for 72 h) were collected with a solid-phase microextraction (SPME) samplers (65 µm polydimethylsiloxane [PDMS] /divinylbenzene [DVB]; Supelco, Inc) by exposing the fibre in the headspace of a glass vial (40 ml) containing 10 ml of the sample for 18 h. Based on preliminary study with multiple fibres (100 µm PDMS, 65 µm PDMS/DVB, 75 µm carboxen/PDMS; Supelco, Inc) with various collection time, the current protocol (i.e., PDMS-DVB for 18 h) for volatile collection was established to maximize the detection sensitivity for the compounds of interest. The volatile collection was conducted in room temperature (25 - 26°C) without agitation of the sample. Immediately following the collection, volatiles absorbed in the SPME fibre were analysed by gas chromatography (GC) – mass spectrometry (MS). Control sample was collected from CW of same amount. For GC-MS, electron impact mass spectra (70 eV) were taken with an Agilent 5975C mass selective detector interfaced to a Agilent 7890A gas chromatograph equipped with a DB-5 column (30 m × 0.32 mm inner diameter, Agilent Technologies). Volatile extracts from SPME fibre were injected in splitless mode, with a temperature program of 50°C for 1 min and then 10°C min^−1^ to 280°C with 5-min hold. The temperature of injector and transfer line was 250 °C. Helium was used as the carrier gas. Six different LW and LW headspace samples prepared with different batches of mosquito larvae were analysed with the same method. Compounds in the samples were identified by comparison of retention times and mass spectra with those of authentic standards. A semi-quantitative estimate of the source concentration for LW volatiles was obtained by comparing the peak integration values from 18h LW headspace collections to those obtained from similar headspace collections from standard solutions (10 ml) containing known concentrations of DMDS, DMTS and sulcatone individually (e.g., 10^−6^ M, 10^−7^ M 10^−8^ M). The ratio of the integration value from the standard preparation that generated the headspace integration value that was most similar to the average integration value of the target compound in LW headspace (i.e., within a range of one order of magnitude) determined the semi-quantitative value of the concentration of that compound in LW. The following equation was used to calculate the source concentration (M) of each target compound in LW.

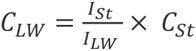

where *C_LW_* is the estimated source concentration of a target compound in LW, *I_St_* is the headspace integration value of the standard preparation closest to the average integration value of the target compound in LW headspace, *I_LW_* is the integration value of the target compound in LW headspace, *C_St_* is the source concentration of standard preparation that generated the headspace integration value that was closest to the average integration value of the target compound in LW headspace.

### Transcuticular electrophysiology

Electroantennogram (EAG), electropalpogram (EPG), and electrolabellogram (ELG) recordings were made from chemosensory organs of two days post blood fed gravid females (subsequently confirmed to contain mature Christopher stage IV or V embryos). EAG assays were carried out one hour after the initiation of scotophase according to previously described methods ^58^, and continued for 3 ~ 4 hours with randomized order of EAG, EPG and ELG. Here, a cold anesthetized gravid female was restrained on slide glass using double sided tape with legs and wings removed as previously described ^32^. The last segment of antenna was subsequently transected and connected to a recording glass electrode filled with Ringer solution (96 mM NaCl, 2 mM KCl, 1 mM MgCl2, 1 mM CaCl2, 5 mM HEPES, pH =7.5) where AgCl coated silver wire was in contact to complete a circuit with reference electrode which was similarly connected into compound eye of the female. The antennal preparation was continuously exposed to humidified, charcoal - filtered air flow (1.84 litre/min) transferred through a borosilicate glass tube (inner diameter = 0.8 cm) using stimulus controller (Syntech, Hilversum, The Netherlands), and the open end of the glass tube was located 5 mm from the antennal preparation. Forty microliters of test or control stimuli were transferred onto a piece of filter paper (10 × 50 mm) which was then placed inside of the Pasteur pipette. LW samples (300 early 3^rd^ or late 4^th^ instars incubated for 72 h) were diluted in ddH_2_O, and test chemical was dissolved and diluted in mineral oil to prepare lower concentrations of test stimuli. Odour was delivered to the antennal preparation for 500 ms through a hole placed on the side of the glass tube located 10 cm from the open end of the tube (1.08 litre/min), and the stimulus odour was mixed with continuous air flow through the hole. A charcoal-filtered air flow (0.76 litre/min) was delivered from another valve through a blank pipette into the glass tube at the same distance from the preparation (i.e., 10 cm from the open end of tube) in order to minimize changes in flow rate during odour stimulation. The test sequence of odours (LW, DMDS, DMTS or sulcatone) was randomized and the order of individual stimuli (i.e., different concentrations of odour and control stimulus) was randomized within each odour session with the interval time of > 40 seconds between every stimulus. Control odours were stimulated before and after the first session of odour (1-octen-3-ol 10^−1^ M for EAG and EPG, oxovaleric acid 10^−1^ M for ELG) in order to check the response sensitivity of test individuals as these compounds have been described as one of most active compounds for each chemosensory organ in previous studies ^32,59^. The resulting signals were amplified 10× and imported into a PC via an intelligent data acquisition controller (IDAC, Syntech, Hilversum, The Netherlands) interface box, and then recordings were analysed by EAG software (EAG Version 2.7, Syntech, Hilversum, The Netherlands). Recordings were replicated on different individual preparations. EPG and ELG recordings and analyses were made using a maxillary palp and labellum of gravid female mosquito following the protocol described above for EAG. In order to enhance the response/noise ratio in EPG and ELG, the tip of palp was not modified except removing mechano-sensory sensilla from the last segment of maxillary palp, and a proboscis was restrained on double-sided tape with labrum removed from labellum. For the analysis of response amplitude, control response (ddH2O or oil) was subtracted from the response amplitude of test stimuli. One sample *t*-test was carried out to determine whether the response amplitude (i.e., control response subtracted) was significantly different from zero (*p* = 0.05, one-sided; JMP 8.01, SAS Institute, Cary, NC). One-sided test was used with the assumption that the polarity of odour stimuli (e.g., depolarization or hyperpolarization) is consistent regardless of concentration. All compounds were obtained from Sigma-Aldrich, Inc. with highest purity available.

### Single sensillum electrophysiological recording

Electrophysiological recordings were conducted on single capitate peg (cp) sensillum along the maxillary palp of female mosquitoes that house three types of olfactory receptor neurons (ORNs) ^32^. Here, gravid females of *An. gambiae* (subsequently confirmed to contain mature Christopher stage IV or V embryos) were cold immobilized (~1 min at -20°C) and mounted on a double-sided tape on a microscope glass slide (25 × 75 × 1.0 mm). Two glass capillaries inserted with chloridized silver wire of appropriate size and filled with 0.1 M KCl saline were used as reference and recording electrode, respectively. The reference electrode was placed in the eye, and the recording electrode was connected to a preamplifier (10×, Syntech, Hilversum, The Netherlands) and inserted at the base of cp sensillum to record the extracellular activity of the ORNs. The signals were digitized by the IDAC4 interface box (Syntech, Hilversum, The Netherlands) and analyzed with Auto Spike v. 3.2 software (Syntech, Hilversum, The Netherlands).

Odour stimuli were diluted in DMSO in decadic steps, ranging from 10^−5^ M to 1 M and DMSO was used as controls. Odour cartridges were prepared by loading filter paper disk (ca. 12.7 mm ø) with 10 µl of test compounds and inserting them into glass Pasteur pipettes connected via silicone tubing to a stimulus controller (Syntech, Hilversum, The Netherlands). Odour stimulation (0.5 litre/min) was carried out for 500 ms by inserting the tip of the SSR odour cartridge into a glass tube with an a charcoal filtered, humidified air-flow (0.5 litre/min) directed towards the maxillary palp which was positioned 10 mm away from end of the glass tube.

The extracellular activity of an individual cp sensillum that houses three physiologically distinct ORNs, cpA, cpB, and cpC was characterized based on the spike amplitudes, spike frequency, and shape as described in figure S5B and a previous study ^32^. The response of an individual neuron to a stimulus was determined by manually counting number of spikes 1000 ms after the onset of neuronal response minus the number of spikes 1000 ms prior to stimulus onset. To rule out the solvent (DMSO) response, solvent spikes were subtracted from the odour induced spike counts. One sample *t*-test (*p* = 0.05, two-sided; JMP 8.01, SAS Institute, Cary, NC) was used to determine whether the normalized response value (i.e., solvent response subtracted) for each neuron was significantly different from zero for each concentration of odour stimuli. If the response value was significantly greater or smaller than zero, the neuronal response was considered as excitatory or inhibitory.

## Acknowledgments

We thank Zhen Li, Samuel A. Ochieng, Emenike O’Kafor, Stephen L. Derryberry, Juan C. Malpartida, Nathaniel T. Day, Emily A. Specht and Alexandra Ruff for mosquito rearing and technical support. We also thank members of the Zwiebel lab for critical reading of the manuscript. This work was conducted with the support of Vanderbilt University and a grant from the National Institutes of Health (NIAID, AI056402) to LJZ. The authors declare no competing interests.

## Authors’ contributions

ES, DHC, AMS and LJZ conceived and designed this study. ES performed the behavioural assays and EAG/EPG/ELG. DHC performed the headspace volatile analyses. AMS performed the SSR. ES, DHC, AMS and LJZ analysed the data. ES, DHC, AMS and LJZ wrote the manuscript.

---

**Figure S1.**
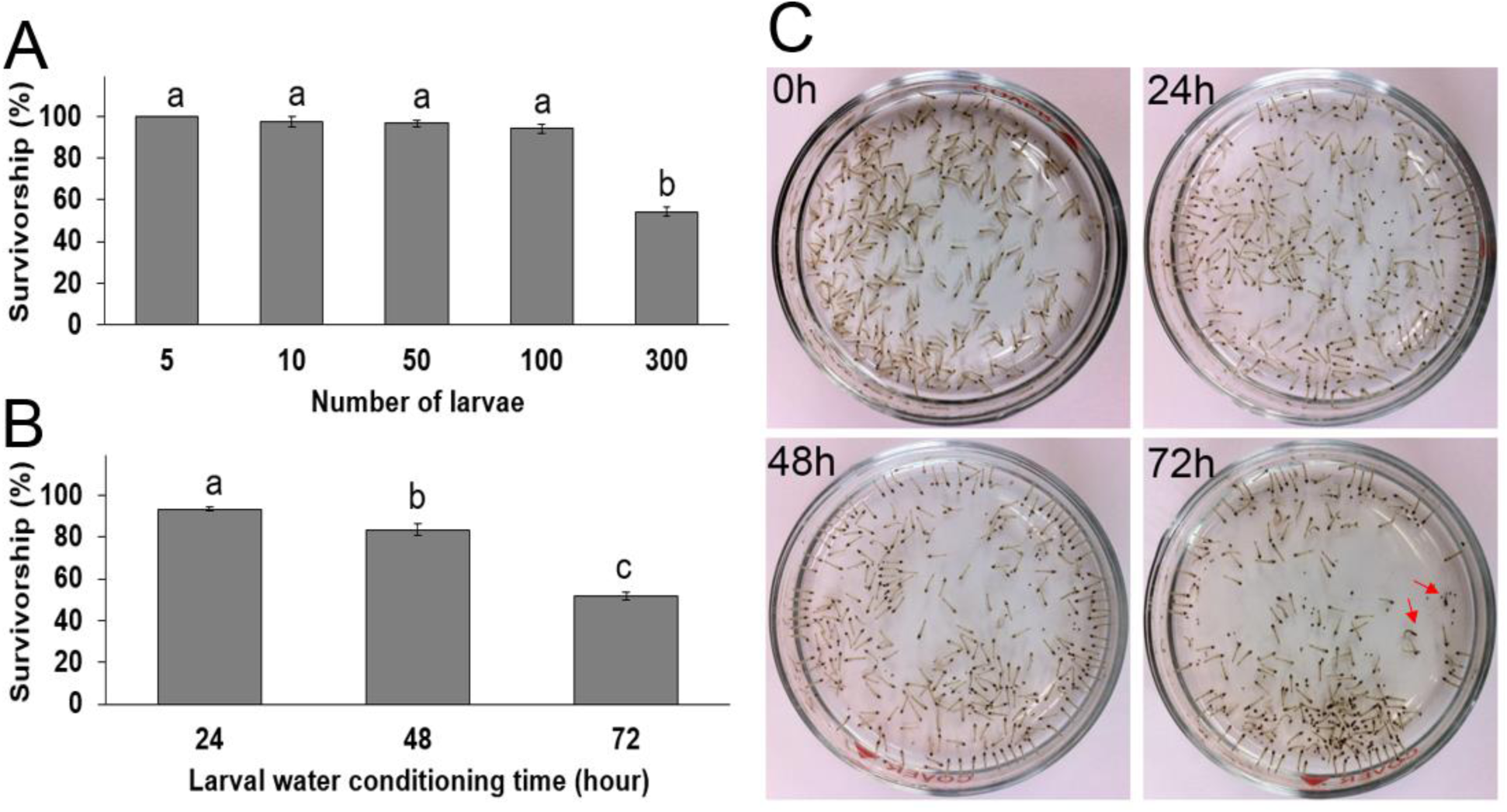
Survival analysis of (A) different number of larvae starved for 72h and (B) 300 larvae starved for differing time period. Differing letters indicate statistical difference at *p* = 0.05 (ANOVA, Tukey post-hoc HSD test). (C) Visual observation of 300 larvae held in 20 ml HPLC water without larval food at four different time points. Red arrows indicate examples of dead larvae at 72h time point.

**Figure S2.**
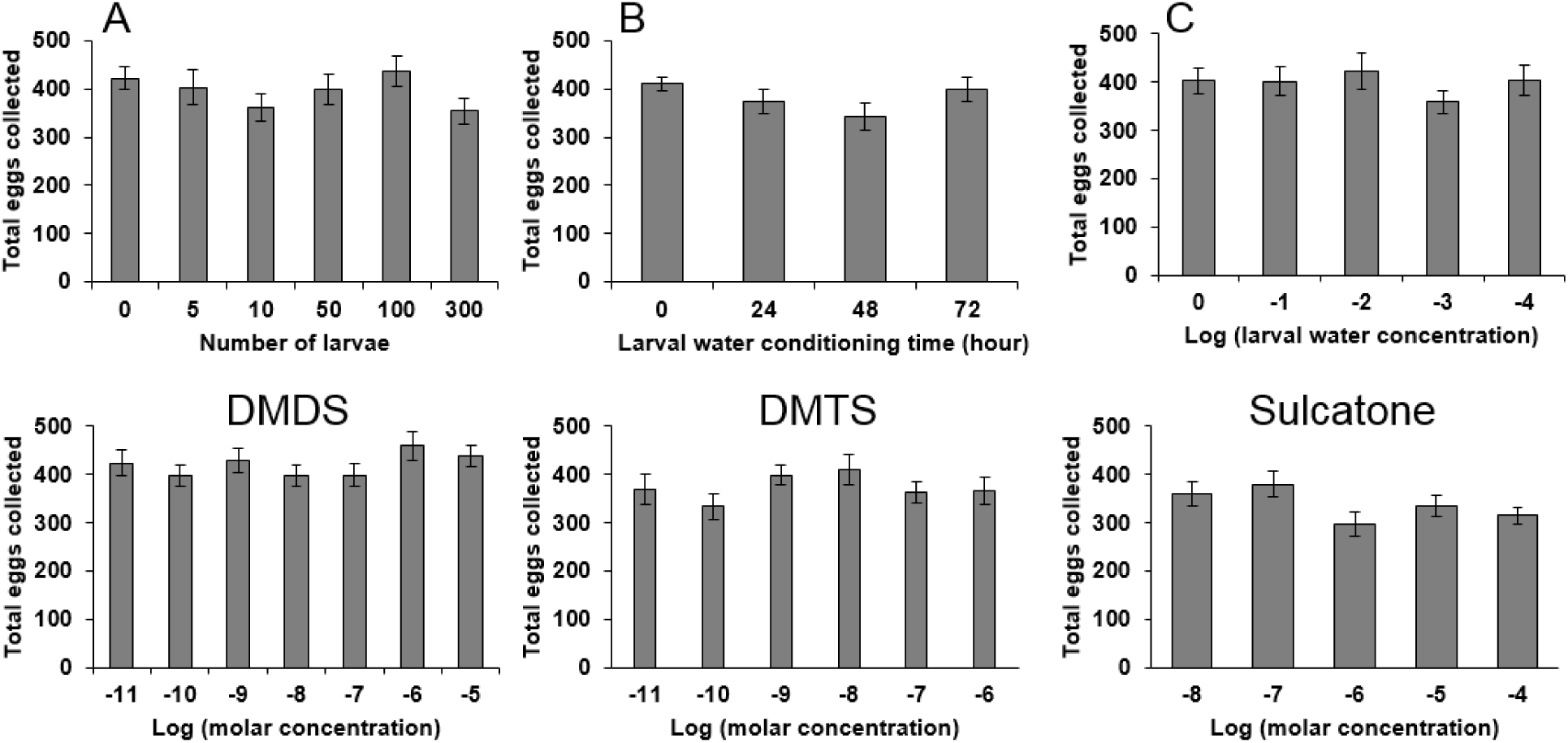
Total number of eggs collected from dual choice oviposition assays. Error bar = s.e.m. Refer number of replicates from figure 1 and 3.

**Figure S3.**
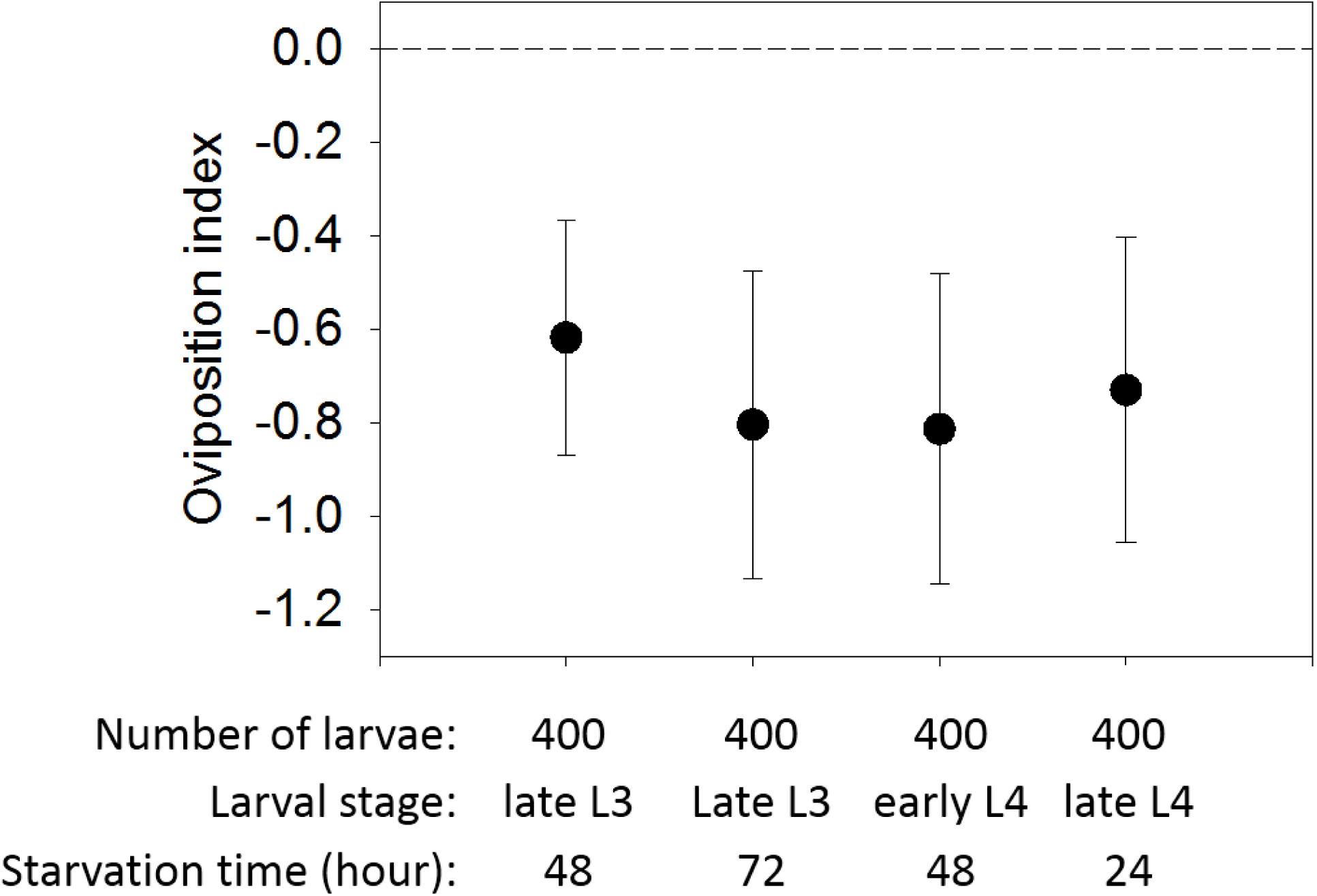
Behavioral response of gravid females of *An. gambiae* in oviposition dual choice assay between control water (CW) and larval water (LW) with varied treatments in number of larvae, age of larvae, and conditioning time. Error bar = s.e.m. (n = 5 ~ 6).

**Figure S4.**
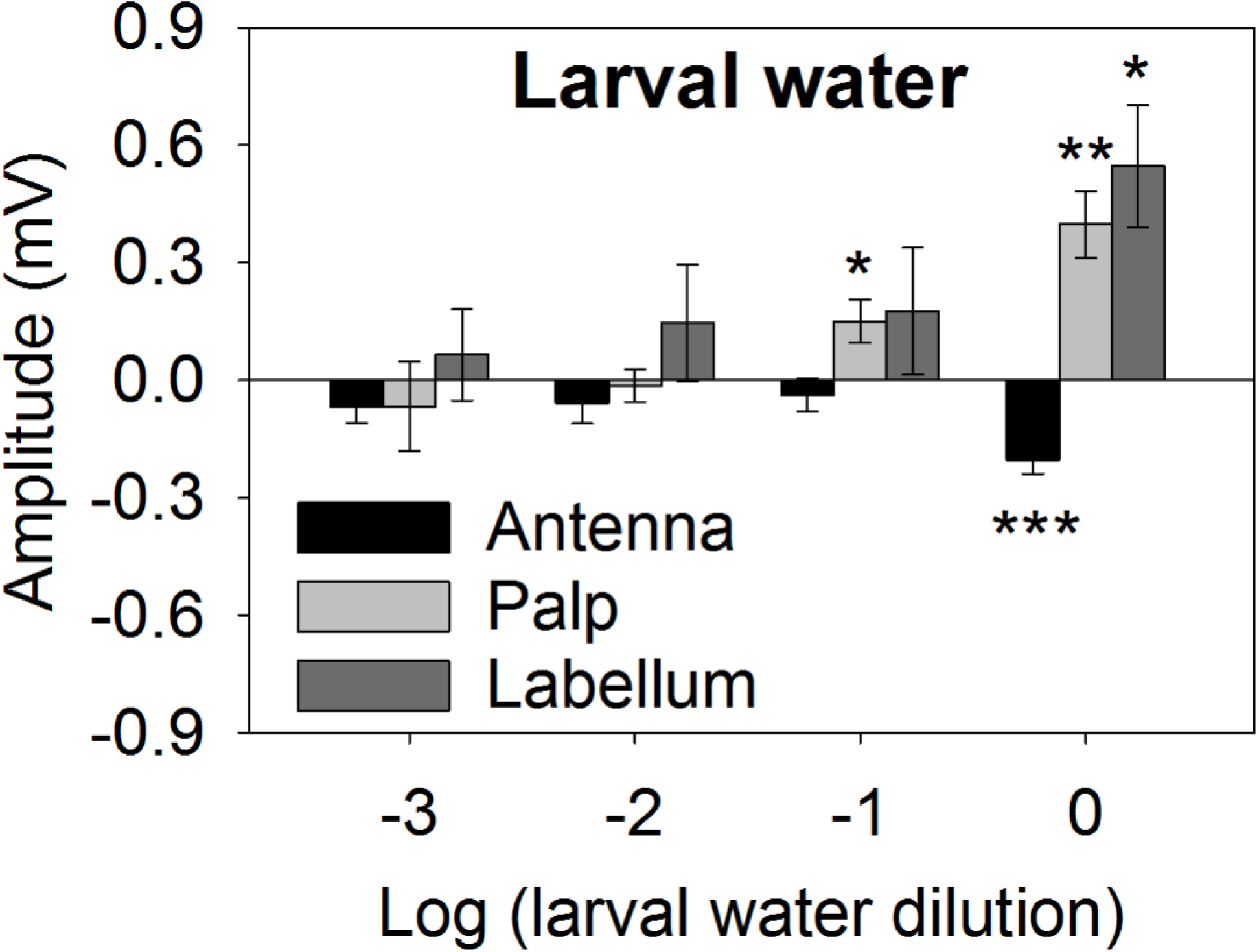
Relative response of antenna, maxillary palp and labellum in EAG/EPG/ELG expressed as response difference to water control (ddH_2_O) of *An. gambiae* females to larval water (300 larvae incubated for 72 h). Y axis represents response amplitude subtracted by control values and X axis represents log transformed larval water dilution. Asterisks represent significant response amplitude different from zero (***, *p* < 0.001; **, *p* < 0.01; *, *p* < 0.05; one sample *t*-test, one-sided). Error bar = s.e.m. (n = 7).

**Figure S5.**
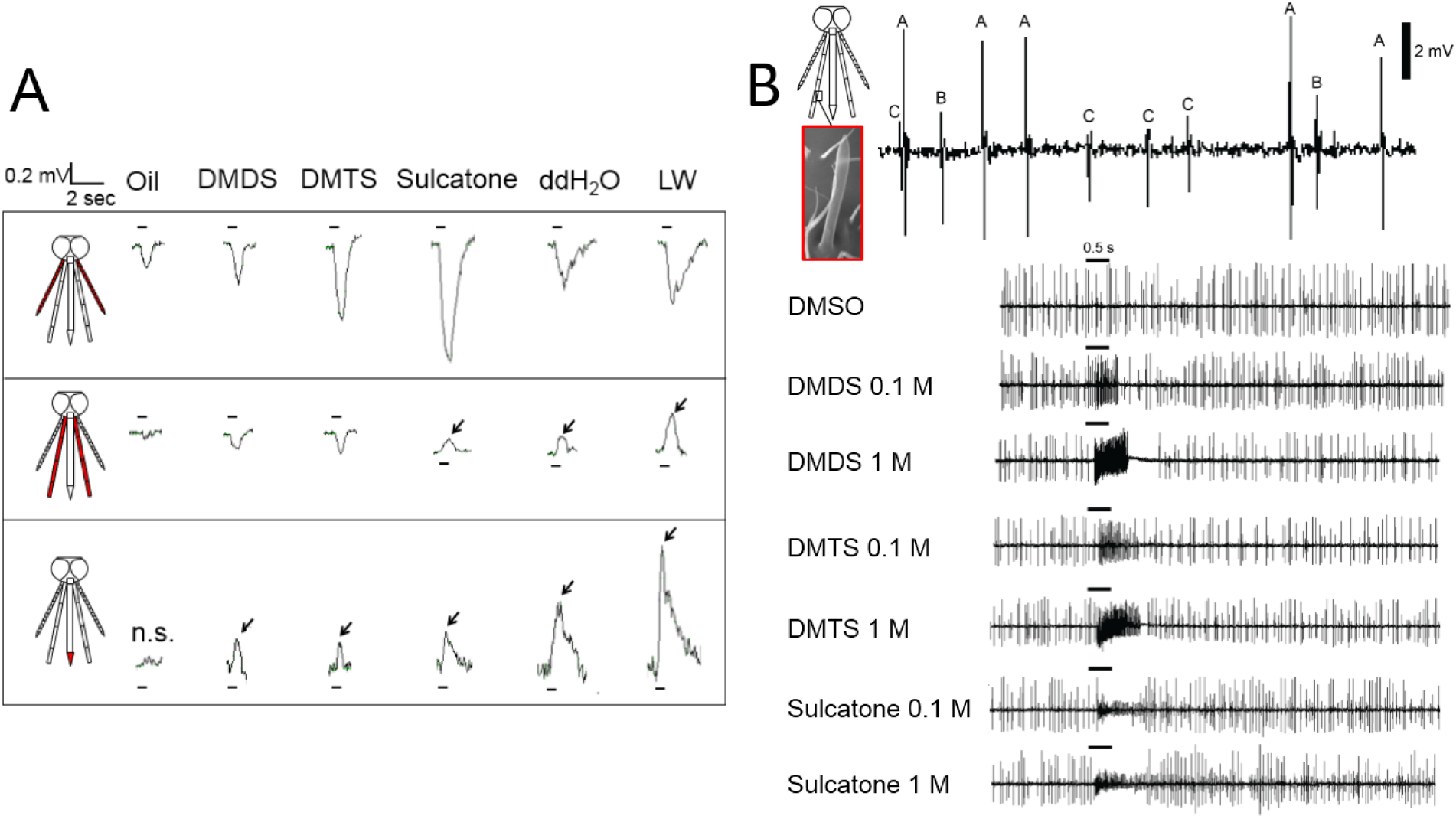
Examples of traces in electrophysiology studies. (A) Differential response kinetics for each odorant (10^−1^ M or undiluted standard larval water) – chemosensory organ combination in EAG/EPG/ELG (top to bottom; each chemosensory organ is highlighted in red in a schematic diagram of mosquito head) and arrows indicate upward responses. (B) Single-sensillum recordings of the responses of the maxillary palp capitate peg sensilla (highlighted in a red box; picture modified from ^5^) of gravid *An. gambiae* females to DMSO, DMDS, DMTS and sulcatone. Action potentials from different neurons are labelled A, B, or C according to spike amplitude and shape. Dark horizontal and vertical bars represent the 500 ms stimulus and 2 mV amplitude, respectively.

